# Genome-engineered *C. elegans* strains with sex-transformed neurons and gut

**DOI:** 10.64898/2026.10.04.756053

**Authors:** Chien-Po Liao, Hongzhu Ji, Oliver Hobert

## Abstract

The sexual identity of specific tissues can be altered through cell-autonomous manipulation of the sex determination pathway in the nematode *Caenorhabditis elegans*. Previously described sexually mosaic strains carried multicopy transgenic arrays that are potentially subject to transgene silencing or misexpression artefacts. We used here the CRISPR/Cas9 system to engineer strains in which coding sequences for either a masculinizing agent (FEM-3) or a feminizing agent (intracellular domain of TRA-2) are inserted with a trans-splicing system at the 3’ end of endogenous genetic loci that are known to be expressed in either a neuron-specific (*rab-3*) or gut-specific (*elt-2*) manner. We validate the tissue-specific sex-transformations induced by these genomic manipulations through expression analysis of known sex-specific genes and through assaying behavioral readouts. These strains may serve the worm community as standard reference tools to study the consequences of tissue-specific sex transformations.

## INTRODUCTION

Sexual dimorphisms are found across all major tissue types throughout the animal kingdom (Bachtrog et al. 2014). In vertebrates, many such dimorphisms are established cell non-autonomously through the action of gonadal hormones (Kashimada and Koopman 2010; Arnold 2017). Many invertebrates, such as *Caenorhabditis elegans*, rely on a cell-autonomous interpretation of sex chromosome configuration to control the expression of sex-specific cellular features (Bachtrog et al. 2014). In *C. elegans*, genetic screens for sex-transformed animals have defined a master regulator of sex identity, the zinc finger transcription factor TRA-1, a Gli family homolog, which promotes hermaphrodite-specific traits at the expense of male-specific traits (Hodgkin 1987; Zarkower and Hodgkin 1992). TRA-1 activity is restricted to hermaphrodites by male-specific degradation of this protein; this degradation is prevented in hermaphrodites by TRA-2 transmembrane receptor signaling and promoted in males by FEM-3 activity (Hodgkin and Brenner 1977; Hodgkin 1987; Kuwabara and Kimble 1995; Schvarzstein and Spence 2006; Starostina et al. 2007). Previous misexpression studies have demonstrated that cell- or tissue-specific overexpression of FEM-3, or expression of a constitutively active intracellular domain of TRA-2 (“TRA-2(ic)”), can reprogram the sexual identity of individual cells. Specifically, cell-specific FEM-3 overexpression cell-autonomously induces male-specific traits in hermaphrodites, while cell-specific TRA-2(ic) expression feminizes cellular traits cell-autonomously in male animals. Some examples of such manipulation are found in (Lee and Portman 2007; White et al. 2007; Mowrey et al. 2014; Ryan et al. 2014; Oren-Suissa et al. 2016; Fagan et al. 2018; Liao et al. 2024).

The expression of the masculinizing factor *fem-3* and the feminizing factor *tra-2(ic)* has traditionally been achieved using multicopy transgenic arrays, either maintained as unstable extrachromosomal elements or as chromosomally integrated arrays. While effective in prior studies (Lee and Portman 2007; White et al. 2007; Mowrey et al. 2014; Ryan et al. 2014; Oren-Suissa et al. 2016; Fagan et al. 2018; Liao et al. 2024), the usage of such transgenes has notable limitations. Extrachromosomal arrays are inherently unstable and mosaic (Stinchcomb et al. 1985; Mello et al. 1991; Mello and Fire 1995) and prone to transgene silencing; the later concern also applying to chromosomally integrated arrays (Kelly et al. 1997; Hsieh et al. 1999). Furthermore, integrated arrays can interfere with genetic mapping by inducing crossover suppression, particularly near closely linked loci of interest. Despite the use of well-characterized promoters to drive tissue-specific expression, multicopy arrays also carry the risk of ectopic expression in unwanted cells. To address these issues, we employed CRISPR/Cas9-mediated genome engineering to insert *fem-3* and *tra-2(ic)* into endogenous loci with known tissue-specific expression. We expect that the resulting alleles that we describe here offer more stable and precise control of sexual identity and may serve as standardized tools for future sex transformation experiments.

## MATERIAL AND METHODS

### *C. elegans* strains and handling

Worms were grown at 20°C on nematode growth medium (NGM) plates seeded with *Escherichia coli* (OP50) bacteria as a food source unless otherwise mentioned. Worms were maintained according to standard protocol. All strains used in this study are listed in **Table S1**.

### CRISPR-Cas9–based genome engineering

We followed an optimized CRISPR/Cas9 protocol for *C*.*elegans* (Eroglu et al. 2023 May 31). To generate *rab-3(ot1401)* and *rab-3 (ot1403)*, crRNA (5’-ttttccctctctcaacatct-3’) and SL2::tra-2(ic) and SL2::fem-3 ssOND were generated by PCR followed by Lamda exonuclease reaction.

#### Sequences for *tra-2(ic)* are

F:5’-

GCGCGTAGCTCTCAGTTTTCCCTCTCTCAACATCTGCTGTCTCATCCTACTTTCACCTAG-3’ and R:5’-TTATAAGTTAATTTACGATCCATATATCTGGTGCCTTCATTAAACCTCTGGGTCTGATAGGTCGC-3’,

#### Sequences for *fem-3* are

F:5’- 5’-

GCGCGTAGCTCTCAGTTTTCCCTCTCTCAACATCTGCTGTCTCATCCTACTTTCACCTAG-3’ and R: 5’-

ATTTATAAGTTAATTTACGATCCATATATCTGGTGCCTTCATCGTTTCCTGGAGCAATCAGTAGC-3’).

To generate *elt-2(ot1438)* and *elt-2(ot1658)*, crRNA (5’- tgcattaaaatggacaactt-3’) and SL2::tra- 2(ic) and SL2::fem-3 ssOND were generated by PCR.

#### Sequences for *tra-2(ic)* are

F:5’-

GATTCTTAAACATATAGAAATATTGCTCAGCTCTTCCAGCTGTCTCATCCTACTTTCA CCTAG-3’ and R: 5’-

GAGAATACAAAACTATGCATTAAAATGGACAACTTTCATTAAACCTCTGGGTCTGATAGGTCGCC-3’ and R: 5’- GATTCTTAAACATATAGAAATATTGCTCAGCTCTTCCAGCTGTCTCATCCTACTTTCA CCTAG-3’

#### Sequences for *fem-3* are

GATTCTTAAACATATAGAAATATTGCTCAGCTCTTCCAGCTGTCTCATCCTACTTTCA CCTAG-3’ and R:5’-

TTTGAGAATACAAAACTATGCATTAAAATGGACAACTTTCATCGTTTCCTGGAGCAA TCAGTAGC-3’).

Because *lin-29* is linked to *rab-3*, we generated *lin-29::SL2::GFP::H2B* in *rab-3(ot1401)* and *rab-3 (ot1403)* using the same reagents as our previous study (Liao et al. 2024).

### Genotyping

To genotype *elt-2(ot1438)*, we used a forward primer (F: 5’- CCAGACTGGCCTAAGACCTGCCG-3’), an outer reverse primer (outR: 5’- GAAACTATACAAGTATGGATGAACAC-3’), and a tra-2(ic)-specific reverse primer (tra-2R: 5’-GATCTAGCATATCTTGAACCAGTTCG-3’). PCR amplification with F and outR were a 429 bp product from the wild-type allele and a 1,767 bp product from the CRISPR/Cas9-edited allele. Amplification with F and tra-2R yielded a 949 bp product only from the CRISPR/Cas9- edited allele.

To genotype *elt-2(ot1658)*, we used a forward primer (F: 5’- CCAGACTGGCCTAAGACCTGCCG-3’), an outer reverse primer (outR: 5’- GAAACTATACAAGTATGGATGAACAC-3’), and a fem-3-specific reverse primer (fem-3R: 5’- CTGAGCTCTCGTTTCTCTGTCAGC -3’). PCR amplification with F and outR were a 429 bp product from the wild-type allele and a 1,847 bp product from the CRISPR/Cas9-edited allele. Amplification with F and fem-3R yielded a 606 bp product only from the CRISPR/Cas9-edited allele.

To genotype *rab-3(ot1401)*, we used a forward primer (F: 5’- GACCCACAGCAACAGCCAAAAGG-3’), an outer reverse primer (outR: 5’- GGGGAATATGATTGAACGTTGGAATAAG-3’), and a tra-2(ic)-specific reverse primer (tra- 2R: 5’-GATCTAGCATATCTTGAACCAGTTCG-3’). PCR amplification with F and outR were a 413 bp product from the wild-type allele and a 1,748 bp product from the CRISPR/Cas9-edited allele. Amplification with F and tra-2R yielded a 893 bp product only from the CRISPR/Cas9- edited allele.

To genotype *rab-3(1403)*, we used a forward primer (F: 5’- GACCCACAGCAACAGCCAAAAGG-3’), an outer reverse primer (outR: 5’- GGGGAATATGATTGAACGTTGGAATAAG-3’), and a fem-3-specific reverse primer (fem- 3R: 5’- CTGAGCTCTCGTTTCTCTGTCAGC -3’). PCR amplification with F and outR were a 413 bp product from the wild-type allele and a 1,828 bp product from the CRISPR/Cas9-edited allele. Amplification with F and fem-3R yielded a 504 bp product only from the CRISPR/Cas9- edited allele.

### Imaging

For the expression analysis of *tra-1(ez72[biotag::GFP::TEV::3xflag::tra-1])* and *lin- 29(ot1664[lin-29::SL2::GFP::H2B]* animals, images were acquired using a 40X with fixed imaging settings on the Zeiss LSM 980 confocal microscope. For the expression of VIT-2::GFP, images of the indicated animals with different backgrounds were acquired using 10X with fixed imaging settings on the Zeiss Axio Imager Z2 compound microscope.

### Locomotion assay

Locomotion was recorded for three minutes and central point movement speed using WormLab (MBF Bioscience, Vermont, USA) (Swierczek et al. 2011).

### Lawn leaving behavior

The assay was performed as previously (Lipton et al. 2004). In brief, assays were conducted on 9 cm NGM plates. 10 µl of concentrated OP50 was placed at the center of each plate, and a single worm was placed on an individual assay plate at the start of the assay. Bacterial tracks created by worm movement were observed at specified time points. An animal was classified as a “leaver” when the track extended 1 cm from the edge of the Petri dish. The index was calculated as (N_total_-N_leaver_)/ N_total_ x 100.

## RESULTS AND DISCUSSION

We fused *fem-3* coding sequences (Ahringer et al. 1992) or the *tra-2ic* coding sequences (Wang and Kimble 2001) behind an SL2-transsplicing intergenic sequence from the *gpd-2/3* operon (Huang & Hirsh, 1989) and inserted these sequencing with the CRISPR/Cas9 system at the 3’ end of the genomic loci of *rab-3* and *elt-2* **(Figure 1A)**, two genes previously reported to be exclusively expressed in neurons and intestine, respectively . Control insertions of an SL2::GFP::H2B cassette reveal the expected GFP signal in the intestine (Leyva-Diaz and Hobert 2022) and gut **(Figure S1)**.

**Figure 1:**
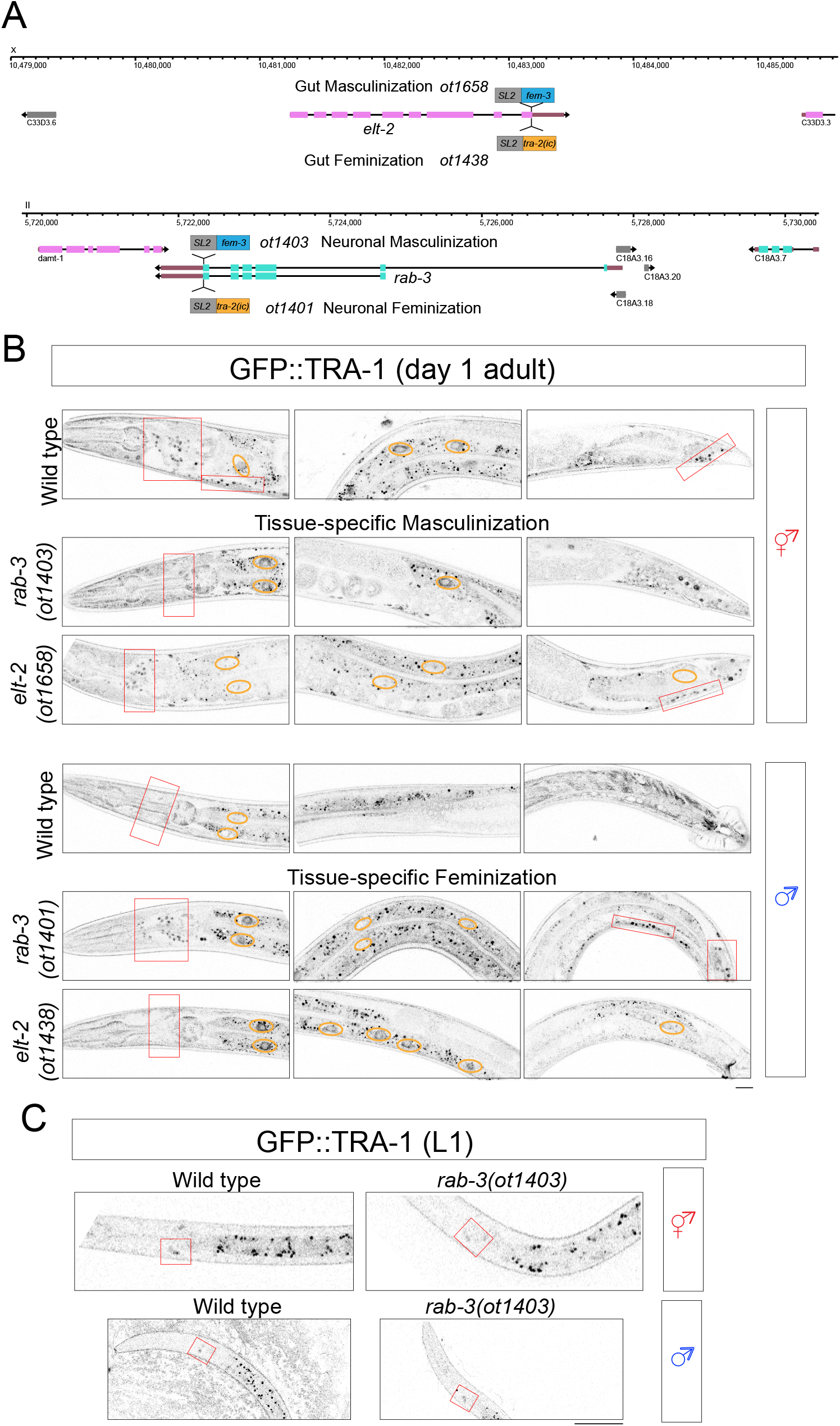
Neuron and gut sex change effect on *tra-1* expression. **(A)** Sex change strains built for this study: *rab-3(ot1403[rab-3::SL2::fem-3])* and *rab-3(ot1401[rab-3::SL2::tra-2ic])* and intestinal sex-change: *elt-2(ot1658[elt-2::SL2::fem-3])* and *elt-2(ot1438[elt-2::SL2::tra-2ic])*. **(B)** *tra-1(ez72[GFP::TEV::3xflag::tra-1])* expression in wild-type, *rab-3(ot1403)*, and *elt-2(ot1658)* hermaphrodites (left), and in wild type, *rab-3(ot1401)*, and *elt-2(ot1438)* males (right) in day 1 young adult. Read boxes indicates location of head neuron ganglia, orange circles indicate gut nuclei. **(C)** *tra-1(ez72[GFP::TEV::3xflag::tra-1])* expression in wild-type, *rab-3(ot1403)* hermaphrodites (left) and in wild type and *rab-3(ot1401)* in L1 stage animals. Scale bar = 10 µm.

We first examined whether endogenous TRA-1 expression is modulated in these strains, as predicted by the tissue-specific genetic manipulation. In wild-type day 1 adults, GFP produced from a *gfp-*tagged *tra-1* locus is broadly expressed in postmitotic cells of hermaphrodites but is largely repressed in males (Bayer et al. 2020). Masculinization of the nervous system in *rab- 3(ot1403[rab-3::SL2::fem-3]* hermaphrodites led to a loss of TRA-1 expression specifically in the nervous system, while expression in other tissues remained unaffected (**Figure 1B**). Similarly, masculinization of the intestine in hermaphrodites in *elt-2(ot1658[elt-2::SL2::fem-3])* animals led to a loss of TRA-1 expression specifically in the intestine, while expression in other tissues remained unaffected **(Figure 1B)**. Conversely, feminization of the intestine in *elt- 2(ot1438[elt-2::SL2::tra-2(ic)])* males resulted in ectopic TRA-1 expression limited to intestinal cells, with no effect on other tissues **(Figure 1B)**. Similarly, feminization of the nervous system in male *rab-3(ot1401[rab-3::SL2::tra-2ic])* animals induced ectopic TRA-1 expression in neurons as predicted, but also unexpectedly in anterior intestinal cells, suggesting a potential cell-nonautonomous regulatory influence of neuronal sexual identity on intestinal TRA-1 expression **(Figure 1B)**.

We assessed the effect of pan-neuronal *fem-3* and *tra-2ic* expression on early larval expression of TRA-1, which is not sex-specific and restricted to two head sensory neuron classes, AWA and ASG (Bayer et al. 2020). We find that *rab-3::SL2::fem-3* does not abolish this expression in either sex, nor is *rab-3::SL2::tra-2ic* capable of inducing TRA-1 expression throughout the nervous system **(Figure 1C)**. These findings indicate that, at early stages, the regulation of TRA-1 expression is distinct from that at later stages, consistent with complex interactions between the sex determination pathway and the heterochronic gene pathway (Berkseth et al. 2013; Bayer et al. 2020).

Returning to fully mature adult animals, we next investigated whether the transcriptional regulation of a downstream TRA-1 target, *lin-29a* (Pereira et al. 2019), aligns with our manipulation of TRA-1 expression. To this end, we utilized a previously generated *lin-29a* SL2- based reporter (Liao et al. 2024). As we previously showed, this transcriptional *lin-29a* reporter is expressed in males in several sex-shared neurons, particularly throughout the ventral nerve cord (VNC), and sparsely in head and tail neurons (Liao et al. 2024), consistent with a translational reporter allele (Pereira et al. 2019). We found that both reporter alleles show male-specific *lin-29a* transcription in intestinal cells of adult animals **(Figure 3** shows the SL2-based reporter allele). Masculinization of neurons or the intestine in *elt-2(ot1658[elt-2::SL2::fem-3])* hermaphrodites induced ectopic *lin-29a* expression in the respective tissues in hermaphrodites, without affecting other cell types **(Figure 2)**. Conversely, feminization of neurons in male *rab- 3(ot1401[rab-3::SL2::tra-2ic])* animals abolished *lin-29a* expression selectively in neurons and feminization of the gut abolished *lin-29a* expression in gut **(Figure 2)**. Notably, despite the ectopic TRA-1 expression observed in the gut of males with feminized neurons (describe above), *lin-29a* expression in the intestine remained unaffected in these animals **(Figure 2)**.

**Figure 2:**
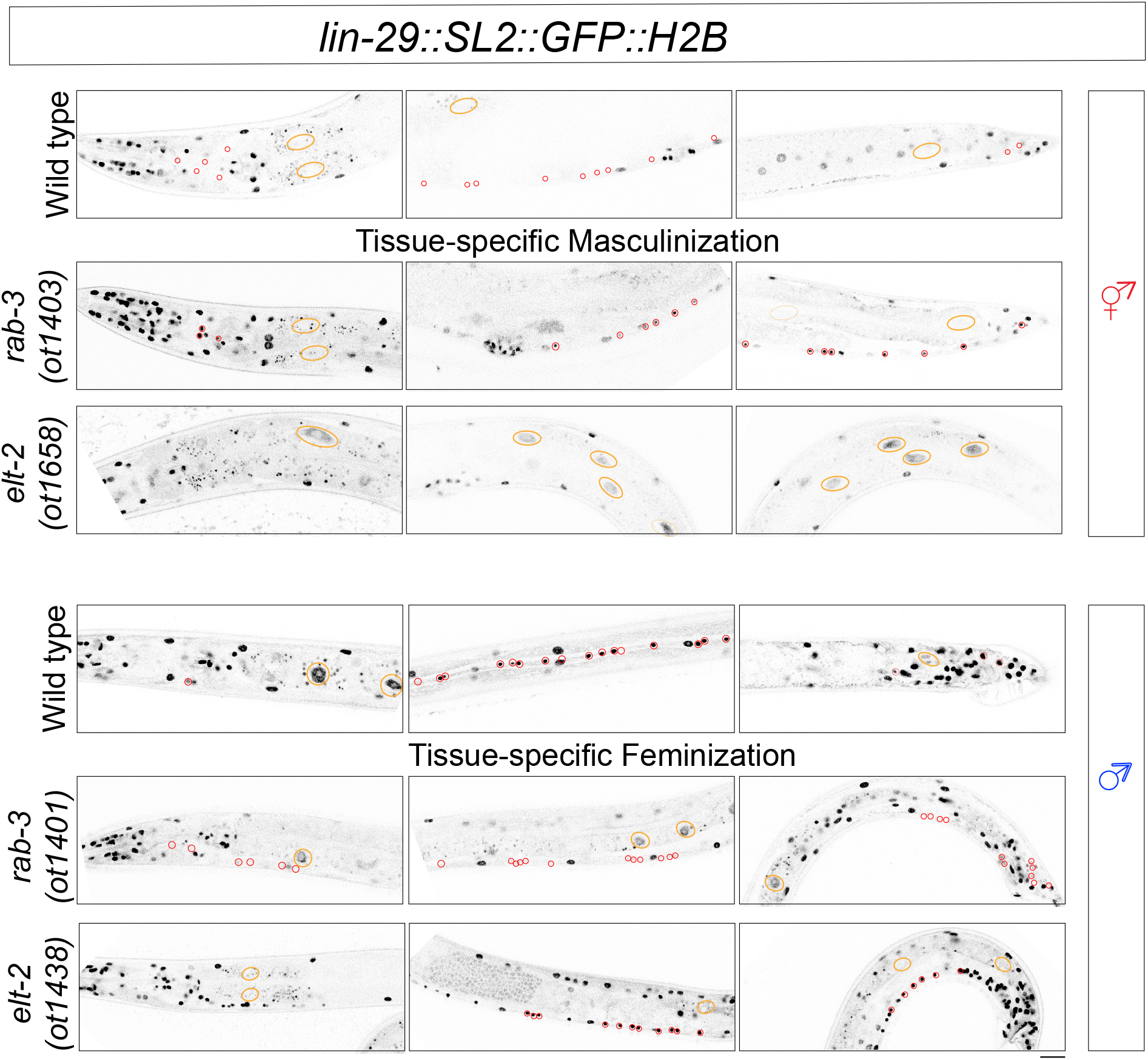
Neuron and gut sex change effect on *lin-29* expression. *lin-29::SL2::H2B::GFP* reporter expression in expression in wild-type, *rab-3(ot1403)*, and *elt-2(ot1658)* hermaphrodites (left), and in wild type, *rab-3(ot1401)*, and *elt-2(ot1438)* males (right). Orange circles indicate intestinal cells, red circles neuronal cels. Scale bar = 10 µm.

**Figure 3:**
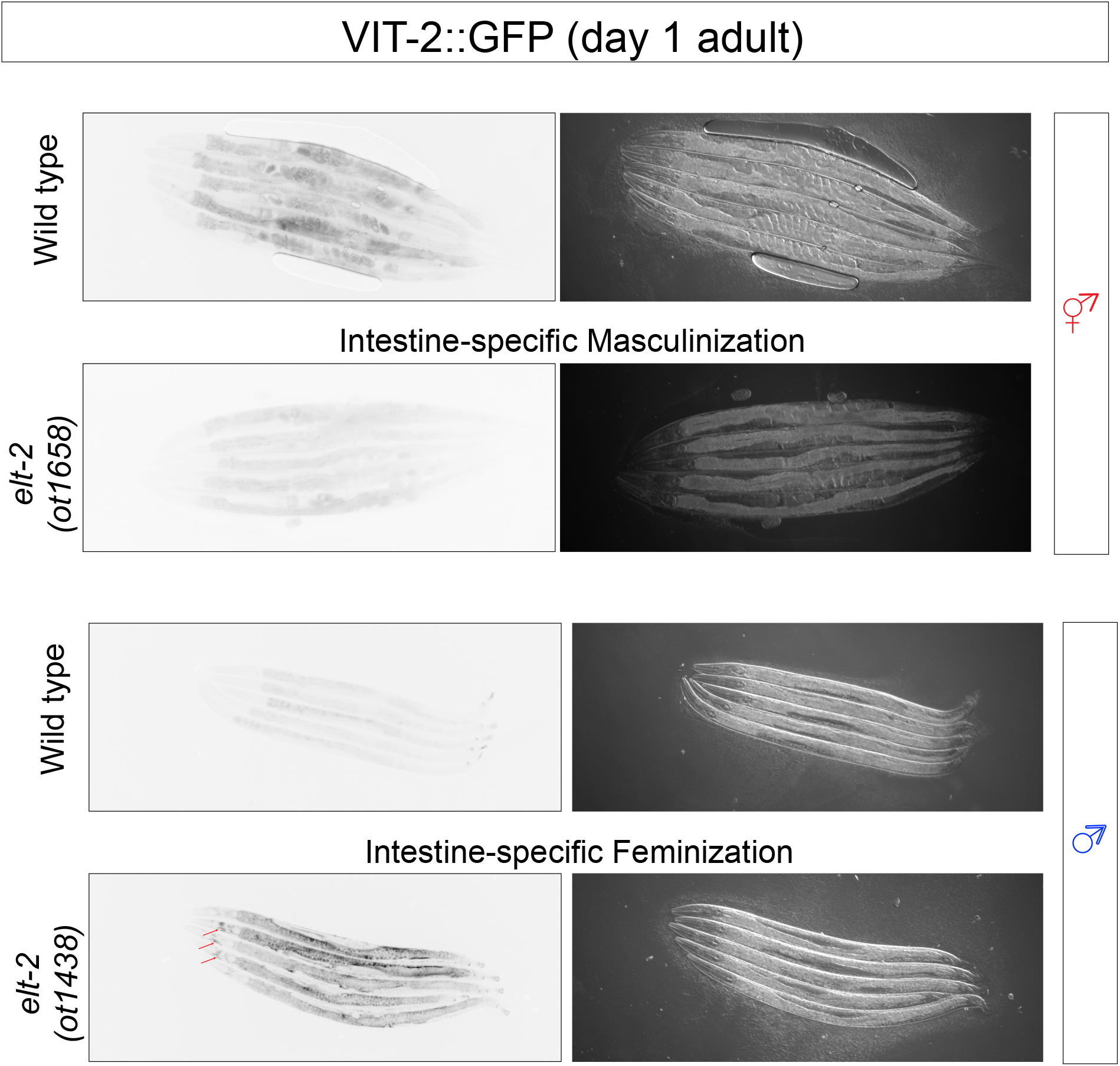
Gut sex change effect on vitellogenin expression. *vit-2(crg9070[vit-2::gfp])* protein expression in wild-type, and *elt-2(ot1658)* hermaphrodites (left) and in wild type, and *elt- 2(ot1438)* males (right) in day 1 young adult.

To further validate the consequences of sex change of the gut in our strains, we examined the endogenous expression of the yolk lipoprotein VIT-2, a vitellogenin produced in the intestine of hermaphrodites and transported to developing oocytes and embryos to support embryogenesis (Spieth et al. 1988; Perez et al. 2017). In males, *vit-2* transcription is repressed by the male-specific DMRT transcription factor MAB-3 (Yi and Zarkower 1999), whose expression in hermaphrodites is directly repressed by TRA-1 (Yi et al. 2000). Using a *vit-2::gfp* reporter allele (Perez et al. 2017), we found that masculinization of the hermaphrodite intestine in *elt- 2(ot1658[elt-2::SL2::fem-3])* animals diminished VIT-2::GFP expression in the intestine (**Figure 3**). In contrast, intestinal feminization induced robust intestinal VIT-2::GFP expression in male animals (**Figure 3)**. Notably, in gut-feminized males, VIT-2::GFP was detected not only in the intestine but also in other tissues. We speculate that this ectopic distribution reflects leakage from the intestine resulting from excessive vitellogenin production **(Figure 3)**.

Finally, we assessed the functional impact of tissue-specific sex identity manipulations. Since behavioral differences between hermaphrodites and males are primarily driven by neuronal sex identity (Mowrey et al. 2014), we analyzed locomotion and exploratory behavior. Using a multi-worm tracker (Swierczek et al. 2011) on a fully seeded bacterial lawn, we observed that wild-type males move faster than hermaphrodites. Masculinization of the nervous system in hermaphrodites increased their movement speed to male-like levels, while feminization of the male nervous system reduced speed to that of hermaphrodites **(Figure 4A)**. Furthermore, males typically exhibit greater exploratory behavior. In lawn-leaving assays (Lipton et al. 2004), feminized males showed reduced exploration, whereas masculinized hermaphrodites displayed enhanced exploratory behavior **(Figure 4B)**.

**Figure 4:**
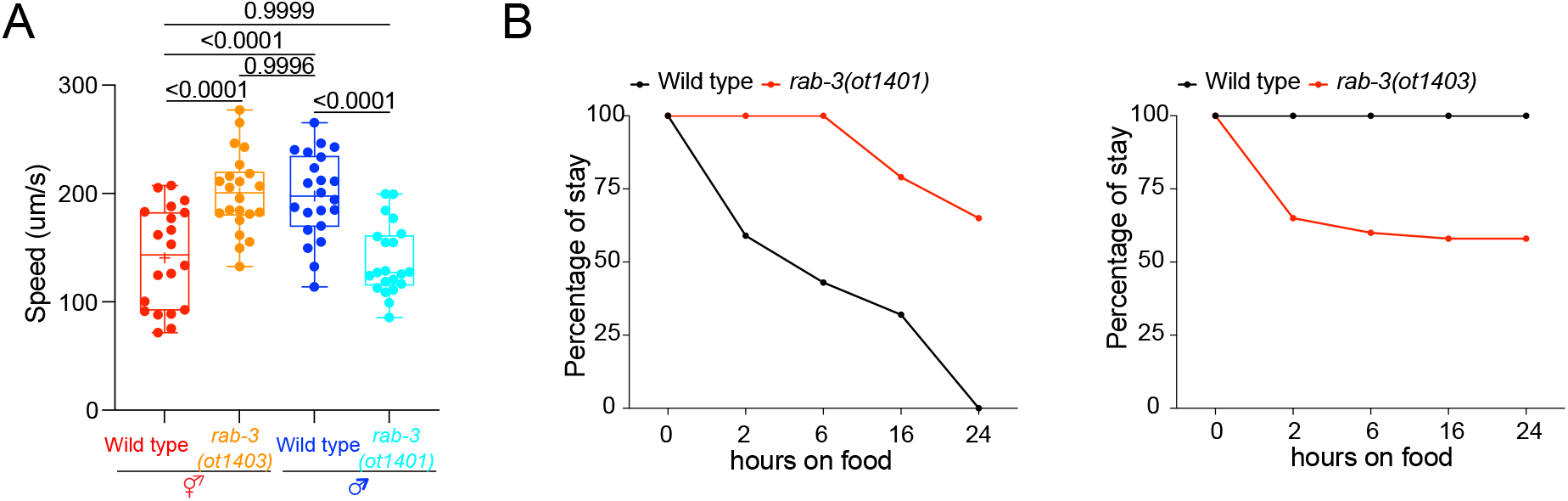
Behavioral analysis of sex-changed strains. **(A)** Quantification of moving speed in wild type and *rab-3(ot1403)* hermaphrodites and wild type and *rab-3(ot1401)* males by multiworm tracker. **(B)** Quantification of the probability of stay in lawn-leaving assay in wild type and *rab- 3(ot1401)* males (left) and wild type and *rab-3(ot1403)* hermaphrodites (right).

Taken together, these results demonstrate that CRISPR/Cas9-mediated engineering of tissue-specific sex identity produces robust and physiologically relevant phenotypes, comparable to those previously achieved using transgenic genetic mani. These strains provide a valuable tool for the community to dissect the roles of sexual identity in the nervous system and the intestine.

## ACKNOWLEDGEMENTS

We thank C. Chen for generating transgenic strains and members of the Hobert Lab for comments the manuscript.

## DATA AVAILABILITY STATEMENT

Strains generated in this paper will be deposited in the Caenorhabditis Genetics Center (CGC).

*SL2::tra-2(ic)* and *SL2::fem-3* sequences can be found in the Supplementary document.

---

**Figure S1:**
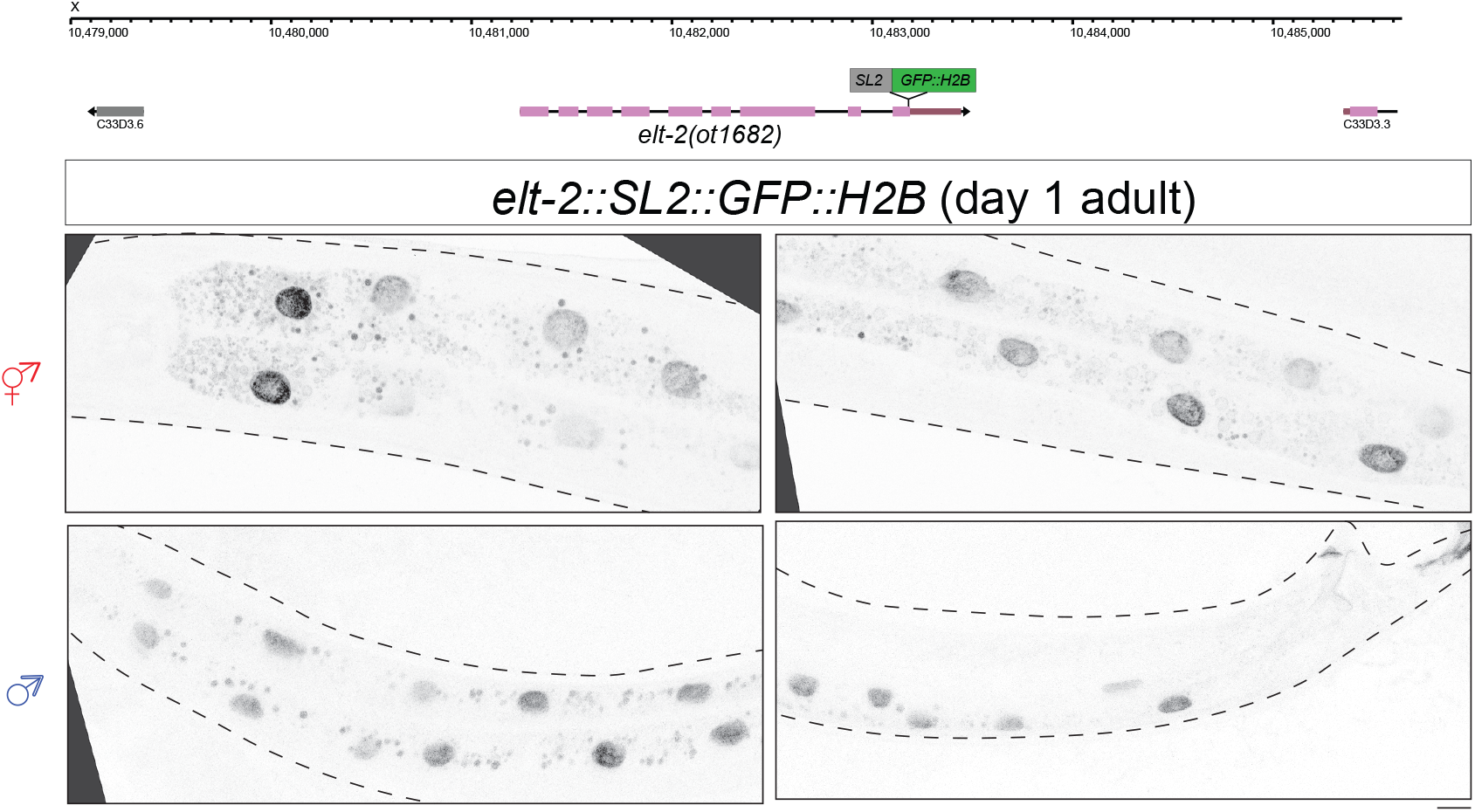
Validation of tissue specificity of SL2-based reporter insertion into the *elt-2* locus. *elt-2(ot1682[elt-2::SL2::H2B::gfp])* is expressed exclusively in intestinal cells in both sexes. Scale bar = 10 µm.

**Table S1:** Strains.

| Strain Name | Genotype |
| --- | --- |
| OH19568 | *rab-3(ot1401[rab-3::SL2::tra-2ic]) II; him-8(e1489) IV* |
| OH18699 | *rab-3(ot1403[rab-3::SL2::fem-3])* |
| OH18882 | *him-8(e1489) IV; elt-2(ot1438[elt-2::SL2::tra-2(ic)]) X* |
| OH19935 | *elt-2(ot1658[elt-2::SL2::fem-3])* |
| OH20037 | *elt-2(ot1682[elt-2::SL2::GFP::H2B])* |
| OH20118 | *tra-1(ez72) III; him-8(e1489) IV* |
| OH20119 | *rab-3(ot1401) II; tra-1(ez72) III; him-8(e1489) IV* |
| OH20120 | *tra-1(ez72) III; him-8(e1489) IV; elt-2 (ot1438) X* |
| OH20121 | *rab-3(ot1403) II; tra-1(ez72) III* |
| OH20122 | *tra-1(ez72) III; elt-2(ot1658) X* |
| OH19114 | *lin-29(ot1482[lin-29::SL2::GFP::H2B]) II; him-5(e1490) V* |
| OH19962 | *rab-3 (ot1401) lin-29(ot1661[lin-29::SL2::GFP::H2B]) II* |
| OH19967 | *rab-3(ot1403) lin-29(ot1664[lin-29::SL2::GFP::H2B]) II* |
| OH20117 | *lin-29(ot1482) II; him-8(e1489) IV; elt-2(ot1438) X* |
| OH19961 | *lin-29(ot1482) II, elt-2(ot1658) X* |
| OH20547 | *dhs-3(ot1691[dhs-3::mScarletI3]) I; him-8(e1489) IV; vit-2(crg9070[vit-2::gfp])X* |
| OH20064 | *dhs-3(ot1691[dhs-3::mScarletI3]) I; him-8(e1489) IV; elt-2(ot1438[elt-2::SL2::tra-2(ic)]) vit-2(crg9070[vit-2::gfp])X* |
| OH20808 | *dhs-3(ot1691[dhs-3::mScarletI3]) I; him-8(e1489) IV; vit-2(crg9070[vit-2::gfp])elt-2(ot1658[elt-2::SL2::fem-3]) X* |

**Table S2:**
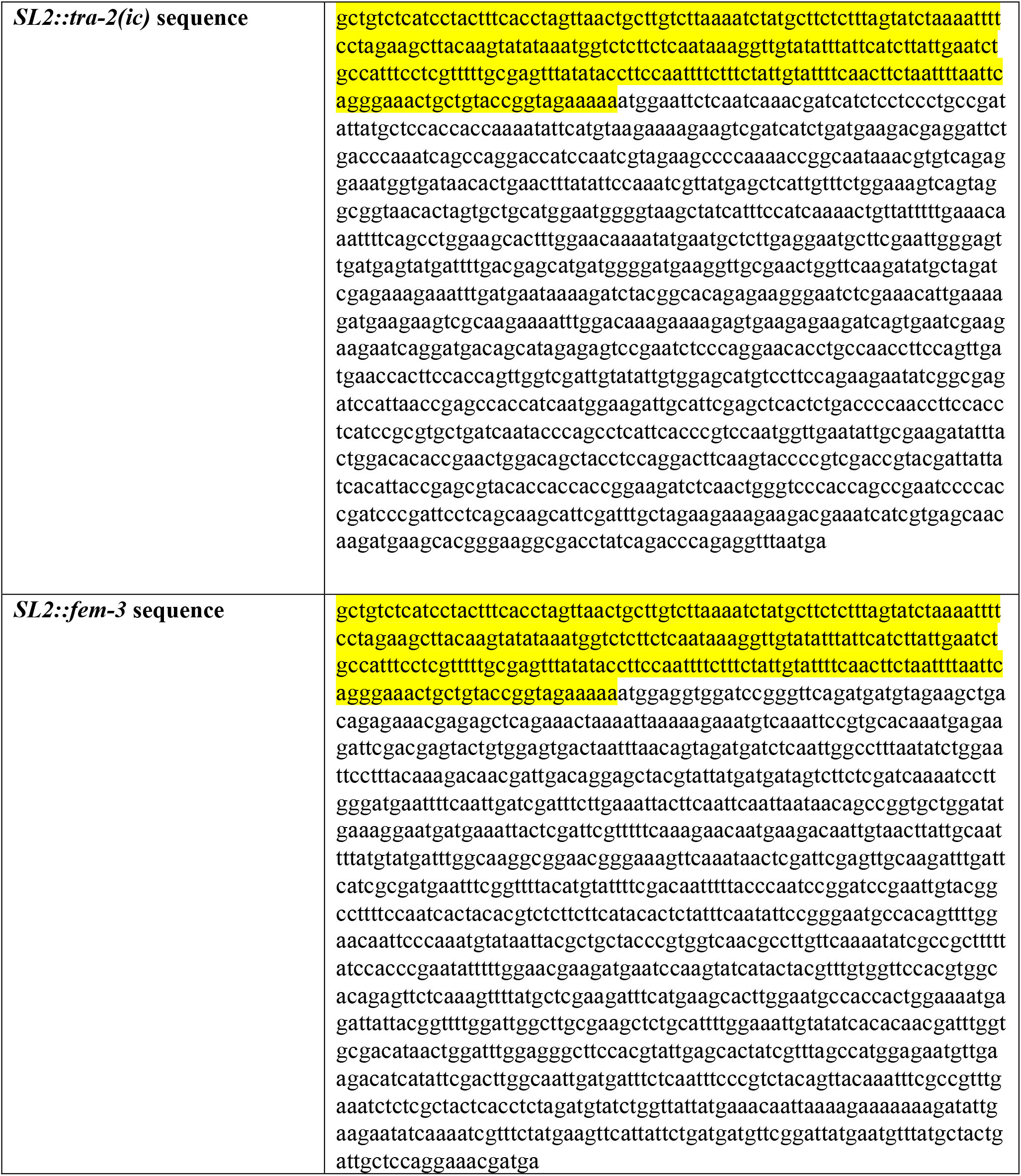
Sequence templates. *SL2* sequence is highlighted in yellow.

